# novoStoic2.0: An integrated framework for pathway synthesis, thermodynamic evaluation, and enzyme selection

**DOI:** 10.1101/2024.09.27.615368

**Authors:** Vikas Upadhyay, Mohit Anand, Costas D. Maranas

## Abstract

Computational pathway design and retro-biosynthetic approaches can facilitate the development of innovative biochemical production routes, biodegradation strategies, and the funneling of multiple precursors into a single bioproduct. However, effective pathway design necessitates a comprehensive understanding of biochemistries, enzyme activities, and thermodynamic feasibility. Herein, we introduce novoStoic2.0, an integrated platform that combines tools for estimating overall stoichiometry, designing de novo synthesis pathways, assessing thermodynamic feasibility, and selecting enzymes. novoStoic2.0 offers a unified web-based interface as a part of the AlphaSynthesis platform (http://novostoic.platform.moleculemaker.org/) tailored for the synthesis of thermodynamically viable pathways as well as the selection of enzymes for re-engineering required for novel reaction steps. We exemplify the utility of the platform to identify novel pathways for hydroxytyrosol synthesis, which are shorter than the known pathways and require reduced cofactor usage. In summary, novoStoic2.0 aims to streamline the process of pathway design contributing to the development of sustainable biotechnological solutions.

## Introduction

Advances in synthetic biology offer considerable potential for engineering biochemical pathways in producing a diverse array of molecules, ranging from biofuels and pharmaceuticals to value-added chemicals and environmentally friendly biodegradation strategies[1–6]. Traditional approaches to biosynthesis often rely on assembling cataloged enzymatic activities, limiting the exploration of novel production routes[7]. Recent studies have demonstrated the utility of leveraging enzyme promiscuity, wherein enzymes exhibit activity on substrates beyond their native targets[8], to enable the assembly of novel pathways through alteration of enzymatic substrate or cofactor specificity. However, the search for novel conversions through enzyme modification presents both a significant enzyme engineering challenge but also an opportunity[9–11]. In an example of a successful application the promiscuous hydroxylase enzyme (4-hydroxyphenylacetate 3-monooxygenase) was used to alter substrate specificity from its native substrate 4-hydroxyphenylacetate to tyrosol and tyramine.[12] These alternative pathways not only minimized the cell metabolic burden by lowering protein synthesis costs but also improved the efficiency of hydroxytyrosol production by rearranging the metabolic flux.

Several pathway design tools are available to generate sequential reaction steps to convert a source chemical into a target molecule. Notable examples include novoStoic[7], RetroPath 2.0[13,14], and BNICE[15], which facilitate the exploration of biochemical pathways. User-friendly web interfaces are provided by tools such as RetroBioCat and novoPathFinder[16,17]. Moreover, recent advancements in machine learning ushered transformer-based models to move beyond the simple molecular-input line-entry system (SMILES) to a Seq2Seq model[18] akin to a large-language model (LLM)[19]. In addition, sampling techniques such as Monte Carlo tree search (MCTS)[18,20] or deep learning-guided AND-OR tree search[20–22] can be used to explore the identification of routes connecting the target molecules with inexpensive precursors.

Although pathway synthesis tools can invoke novel steps to complete biosynthetic pathways, the discovery or re-design of an enzyme to carry out the hypothesized transformation remains a challenge[23–25]. Recently, we developed EnzRank[23], which relies on utilizing convolutional neural networks (CNNs) to understand underlying residue patterns and combines with the substrate molecule signature to provide a probability score for the compatibility of enzyme-substrate pairs. However, implementing the novel steps requires *de novo* enzyme design or protein re-engineering to alter substrate specificity using directed evolution. One such example is the design of a luciferase enzyme that selectively catalyzes the oxidative chemiluminescence of novel luciferin substrates diphenylterazine and 2-deoxycoelenterazine[26].

Thermodynamic feasibility assessment of the entire pathway and also of individual steps is an important check as most databases, used to train ML retrosynthesis tools, treat reactions as reversible, resulting in erroneously adding steps in a thermodynamically unfavorable direction. Tools such as eQuilibrator[27] and dGPredictor[28] can estimate standard Gibbs energy change of reactions. While eQuilibrator uses an expert-defined functional group, dGPredictor uses automated chemical moieties that classify every atom in a molecule based on their surrounding atoms and bonds to feature molecules. Using structure-agnostic chemical moieties in dGPredictor allows for the estimation of standard Gibbs energy of reactions containing novel metabolites absent from databases or molecular structures that cannot be decomposed using expert-defined functional groups.

Herein, we introduce the novoStoic2.0 framework, which integrates the aforementioned tasks (Fig 1) within a single interface. First, the optStoic[29] tool can be used to estimate the optimal overall stoichiometry of the desired conversion by maximizing the yield of the target molecule from the given starting compound. Next, novoStoic[7] attempts to identify the link between the input and output molecules of the overall conversion using both database and novel reactions. To assess the thermodynamic feasibility of these individual steps, including novel ones, dGPredictor[28] is accessed to estimate the standard Gibbs energy changes. Finally, EnzRank[9] can be used to select enzyme candidates for (any) novel conversions identified within the pathway design.

**Fig 1:**
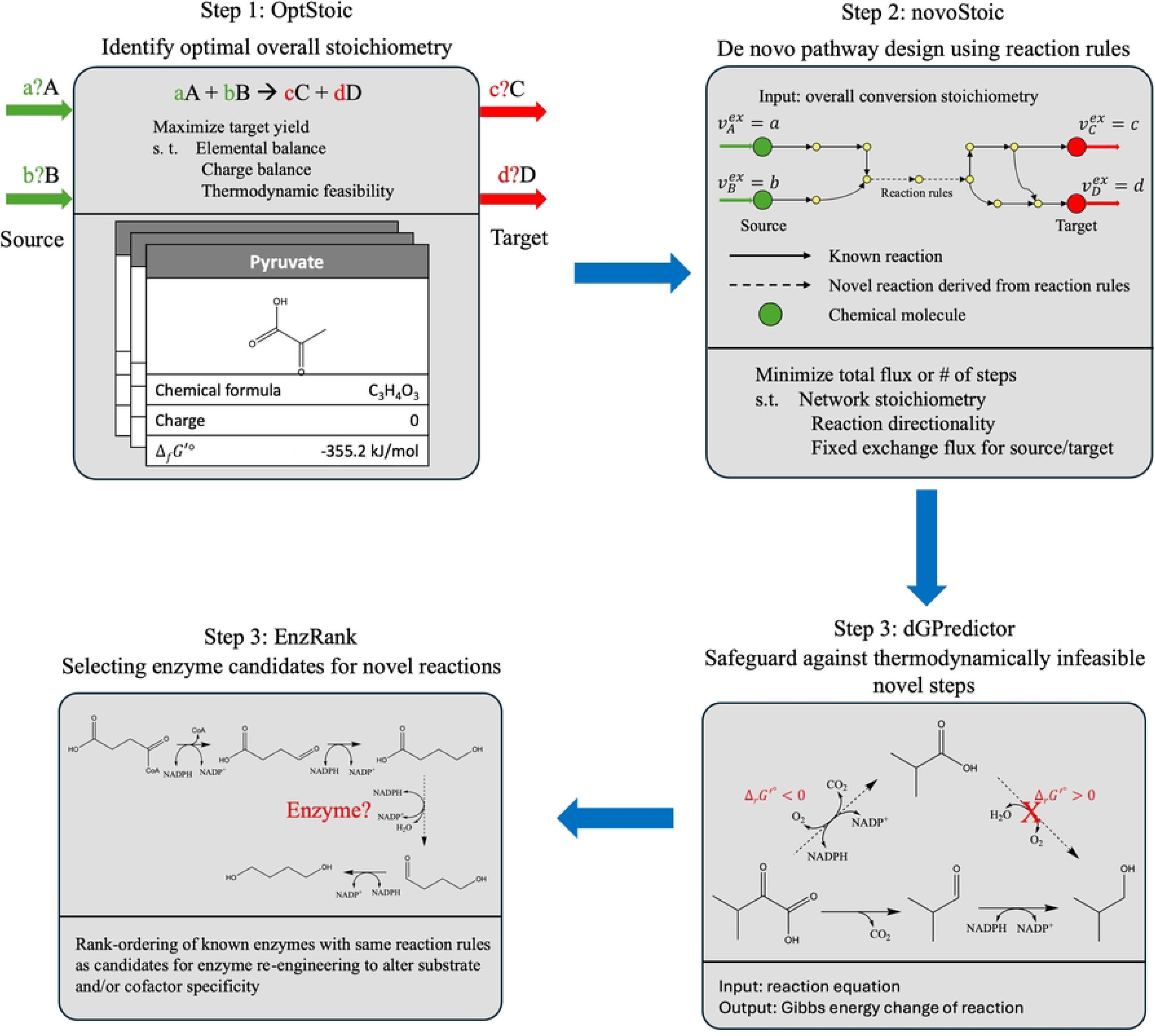
novoStoic2.0: overall workflow for the integrated pathway design including tools for optimal overall stoichiometry of conversion, optStoic. De novo pathway design using optimal stoichiometry, novoStoic. Thermodynamic assessment of reaction steps using dGPredictor and rank-ordering of known enzymes for novel substrate activity to identify starting enzymes for novel steps using EnzRank.

novoStoic2.0 is a one-stop web-based interface for designing biosynthesis pathways that are not only thermodynamically feasible and carbon/energy balanced but also provide suggestions on enzyme selection for re-engineering associated with novel reaction steps.

## Results

### novoStoic2.0 user interface

Fig 2 illustrates the homepage of the web-based platform built using the Streamlit Python package.

**Fig 2:**
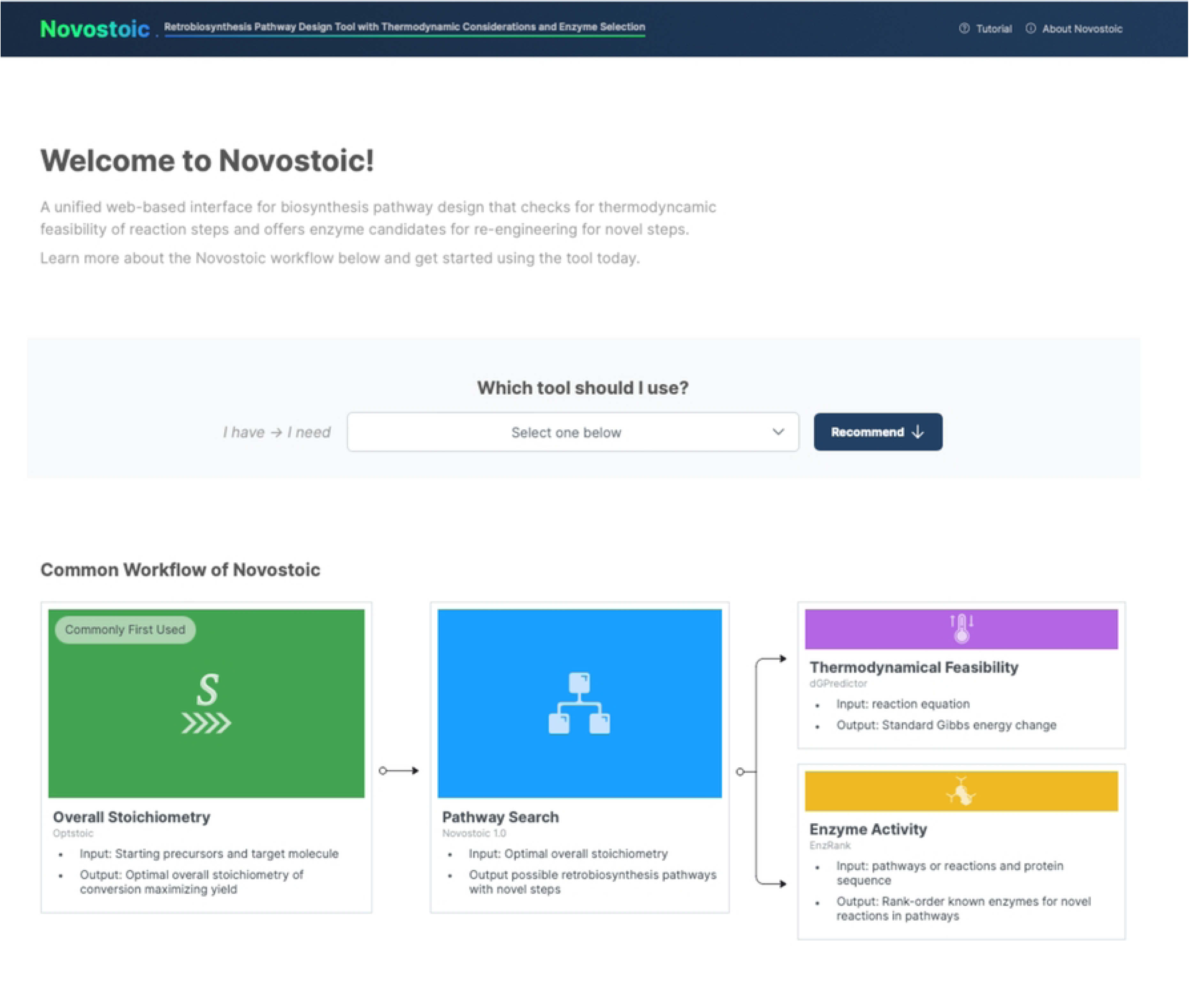
Screenshot of the home page of the novoStoic2.0 user interface. This page provides an option for user to selection the options to choose the specific tool alone as well as in the given workflow to include all the tools for designing pathways.

It provides brief information for each tool along with relevant references. optStoic[29] uses MetaNetX[30,31] as well as KEGG database compound IDs to input both the source and target molecules. Users may also specify any other co-substrate and co-product to be considered in conjunction with the source and target molecules. Subsequently, optStoic (interface shown in Fig 3) solves an LP optimization problem, generally aiming to maximize theoretical yield while ensuring mass, energy, charge, and atom balance[29]. The resultant output of this step is the overall target stoichiometry of the conversion.

**Fig 3:**
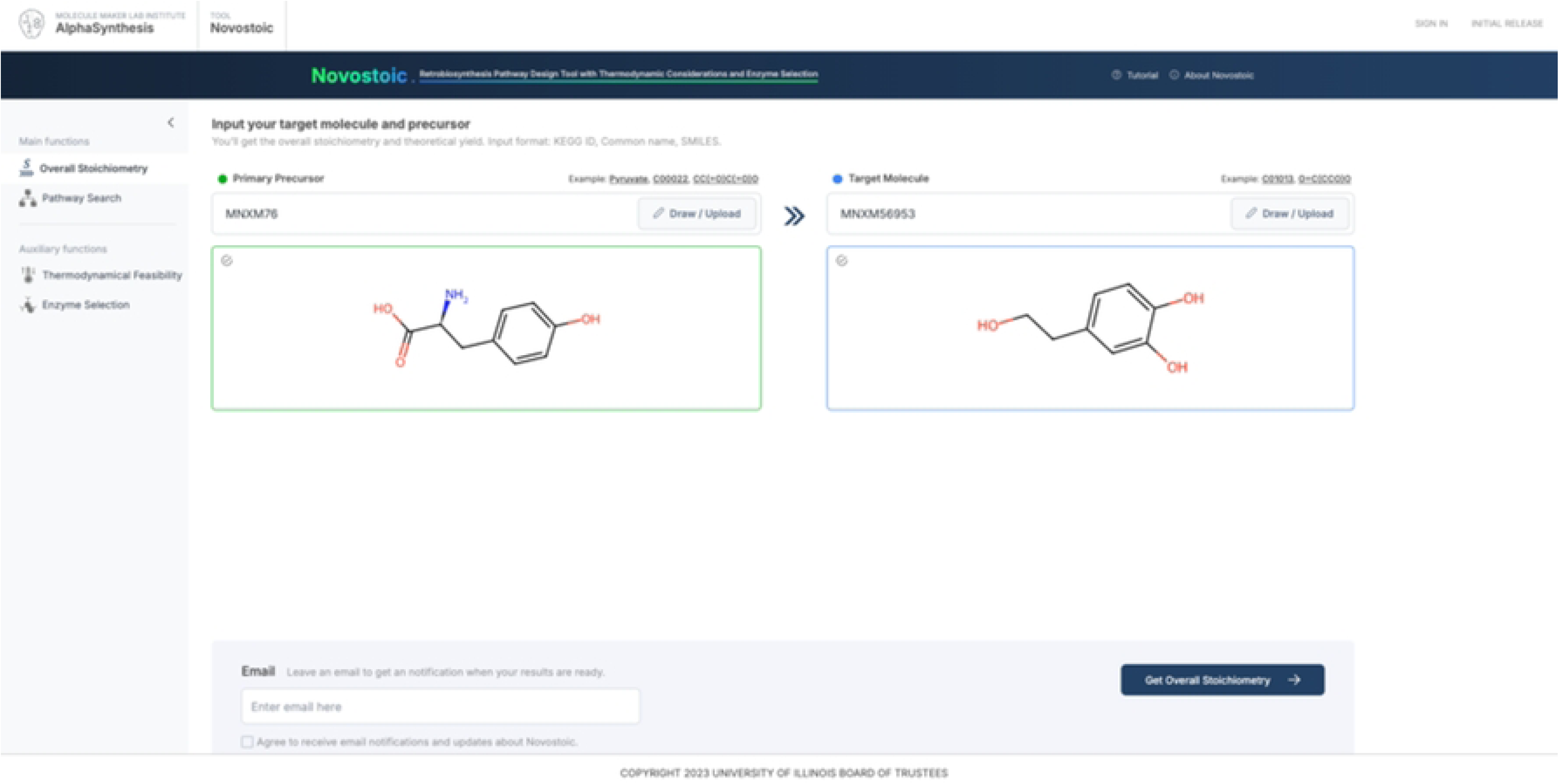
Screenshot of the user interface of optStoic. optStoic requires the MetaNetX ID of the starting and target molecules and outputs the optimal overall conversion stoichiometry, that is used in novoStoic to find synthesis pathways.

This overall stoichiometry becomes a key input for novoStoic, depicted in Fig 4. novoStoic requires additional inputs, including the maximum number of (novel) steps, the maximum number of pathway designs, and the primary source and target molecules.

**Fig 4:**
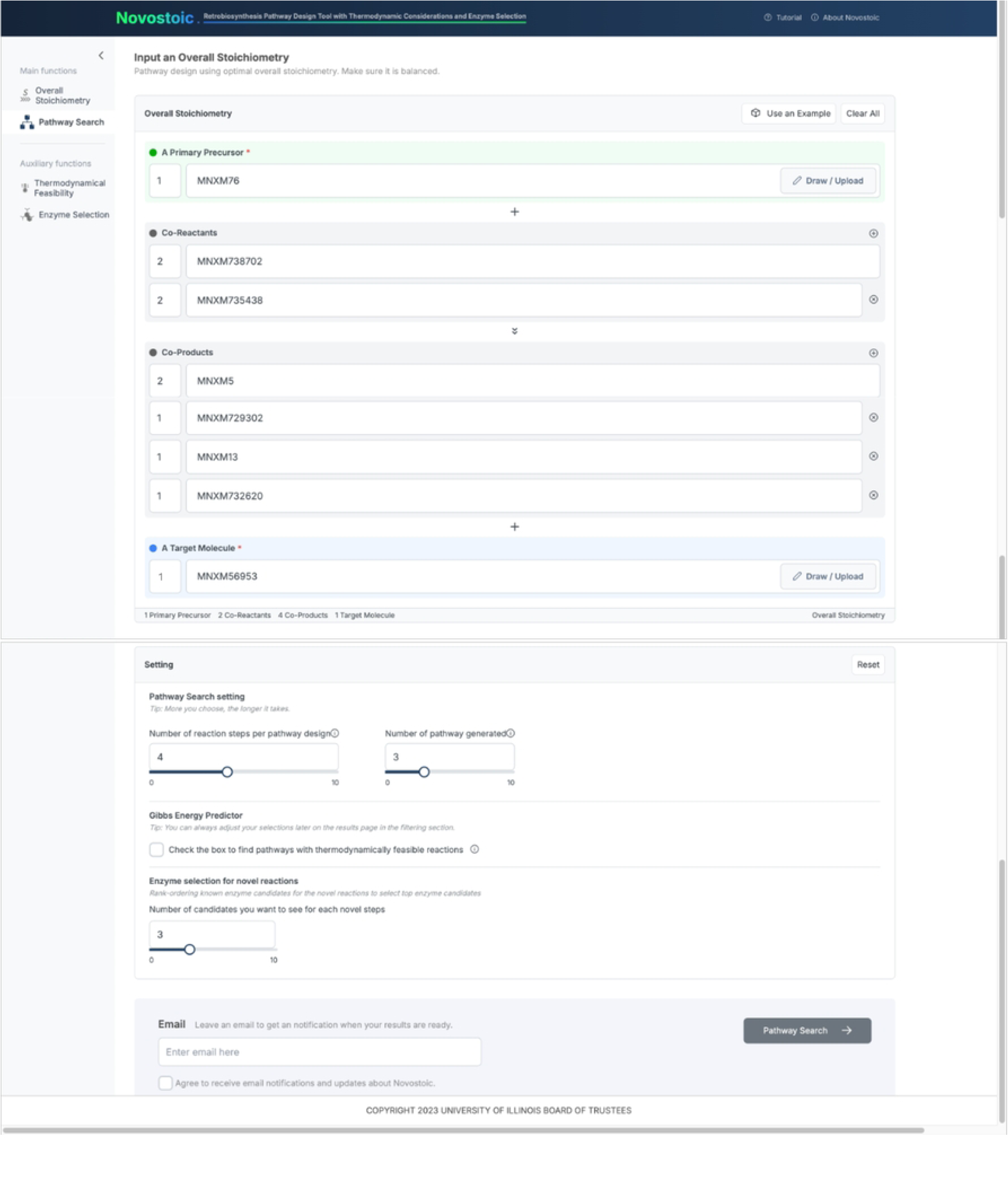
Screenshot of the user interface of novoStoic, which requires overall balanced stoichiometry of conversion from a source molecule to a target molecule, as well as parameters such as maximum number of steps for pathway search as well as number of pathways to search for. For novel steps identified in the pathways, another parameter is required for the number of top known enzyme candidates ranked by EnzRank.

The output of novoStoic visualizes the identified pathways, as well as individual reactions with their standard Gibbs energy estimated using dGPredictor[28]. For pathways involving novel steps, the output provides the option of using EnzRank to generate the top known enzyme candidates for potential activity with novel substrate (i.e., the default is the top five enzyme candidates for each novel step).

dGPredictor could also be used as a standalone standard Gibbs energy estimator as illustrated in Fig 5 using either the KEGG[31] reaction string or the InChI string for novel molecules, that are not present in the KEGG[31] database. This feature holds broad applicability, including conducting global thermodynamic feasibility analysis of genome-scale metabolic models[15].

**Fig 5:**
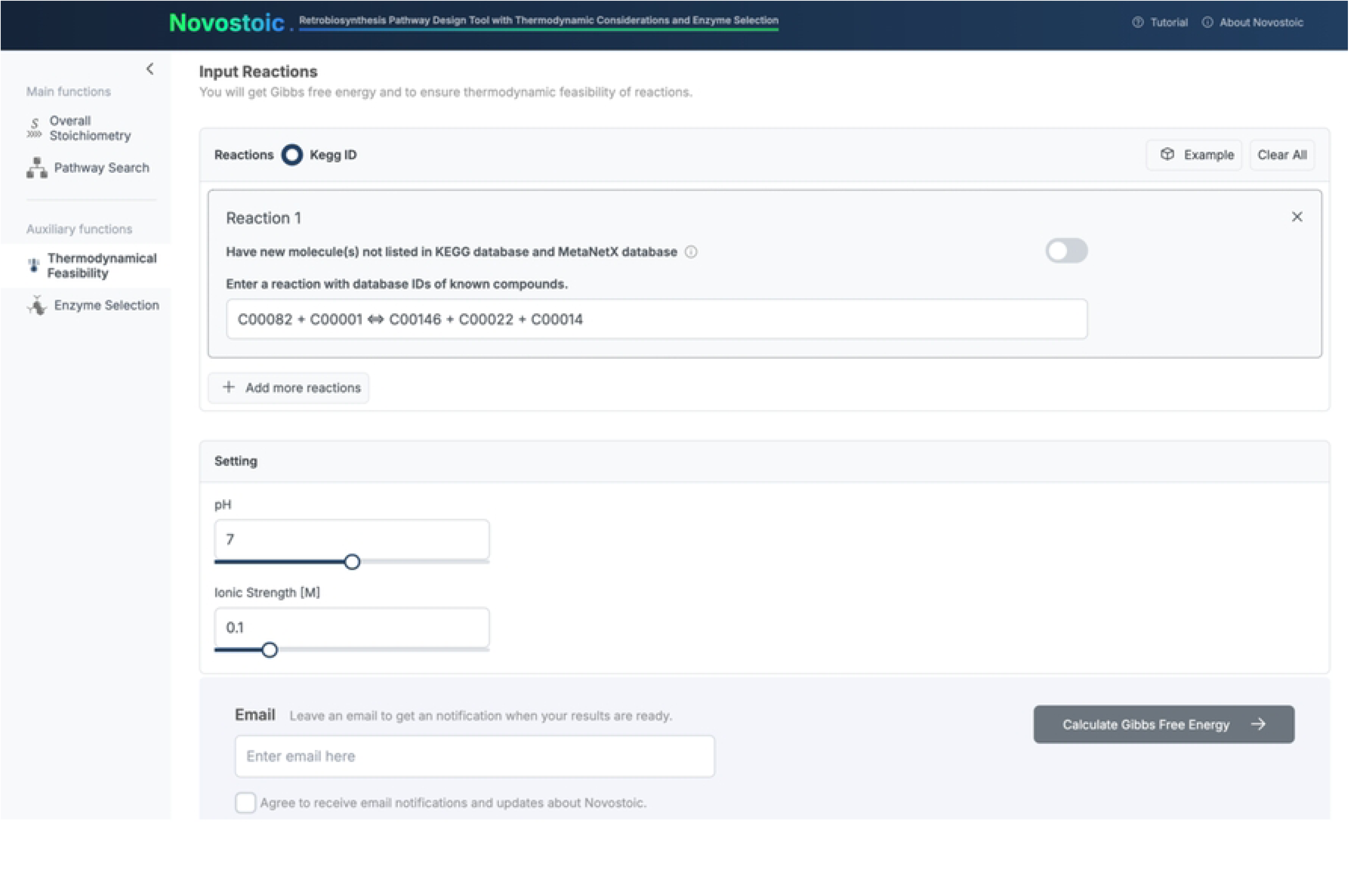
Screenshot of the user interface for EnzRank within novoStoic2.0, requires the SMILES string of the substrate and the enzyme sequence as input to estimate the probability score for the enzyme to have an activity on the substrate.

Finally, as shown in Fig 6, we also provide the user interface of EnzRank as a standalone tool within novoStoic2.0. Here, the users can input the enzyme’s amino acid sequence and the KEGG id or SMILES string (if there is a novel structure) of the substrate to generate the probability score for the enzyme-substrate activity. This score can rank-order known enzymes for novel substrate activity.

**Fig 6:**
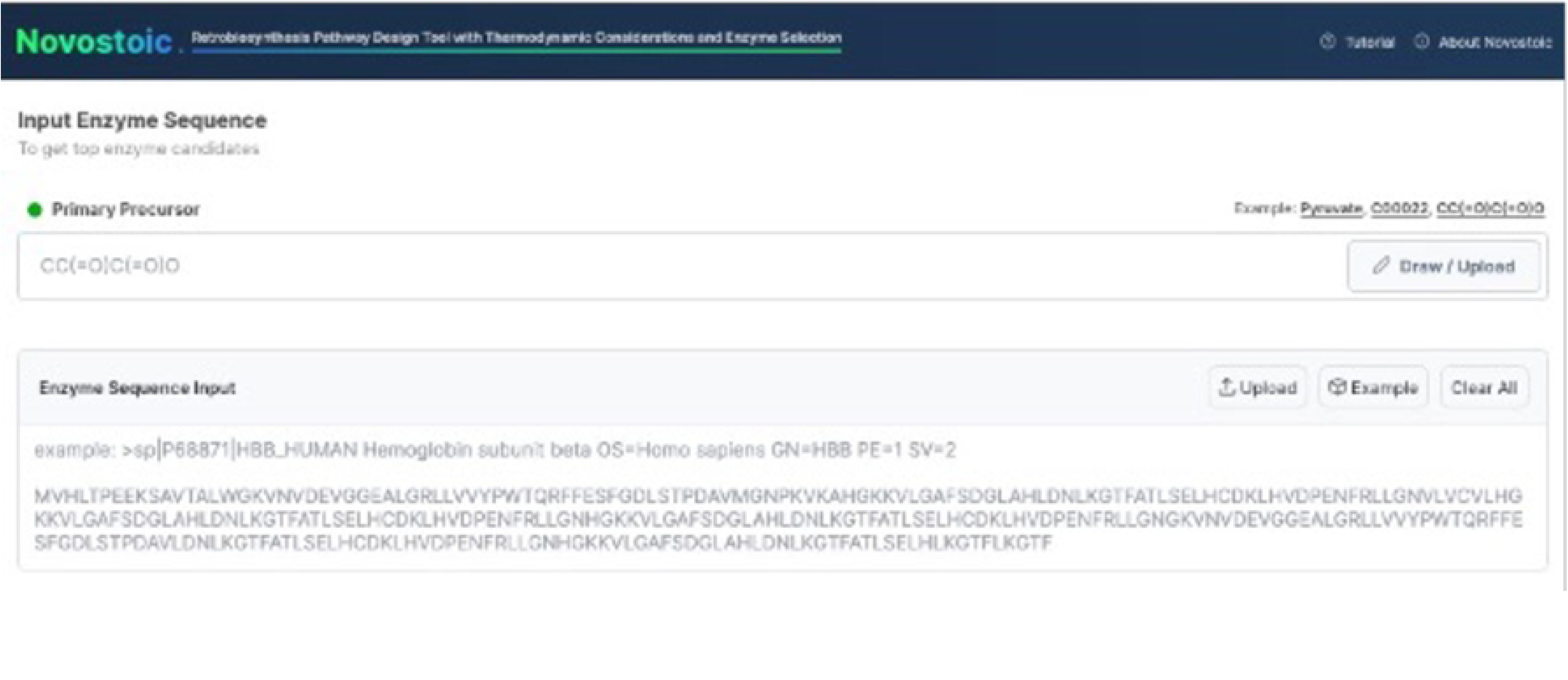
Screenshot for the user interface of dGPredictor within novoStoic2.0, which requires the KEGG reaction string as input. If a molecule is not present in the database, an InChI string is required.

### Example application: Hydroxytyrosol synthesis

Hydroxytyrosol (HXT) is a powerful antioxidant that offers protection from free radicals. The initial effort of biosynthesis uses tyrosine as a starting substrate using tyrosol as the intermediate was reported to achieve a yield close to 50%[12]. A recent effort by Chen et. al[12] uses a promiscuous hydroxylase enzyme to identify two novel pathways for HXT biosynthesis. Motivated by these results herein, we applied novoStoic2.0 to explore additional pathway designs. We show two of the identified new pathways designed using L-tyrosine as a starting precursor and HXT as the target molecule.

The first step involves the identification of the optimal stoichiometry (i.e., max carbon yield) for the overall conversion using optStoic. We explored multiple alternatives in one run by allowing as reactants/products an inclusive set of small molecules CO_2_, H_2_O, NH ^+^, O_2_, H_2_O_2,_ and cofactors such as NAD(P)H, NAD(P)^+^ to balance the overall conversion.

optStoic identified the overall stoichiometry associated with the pathway identified by Chen et. al that uses a promiscuous hydroxylase enzyme given by:

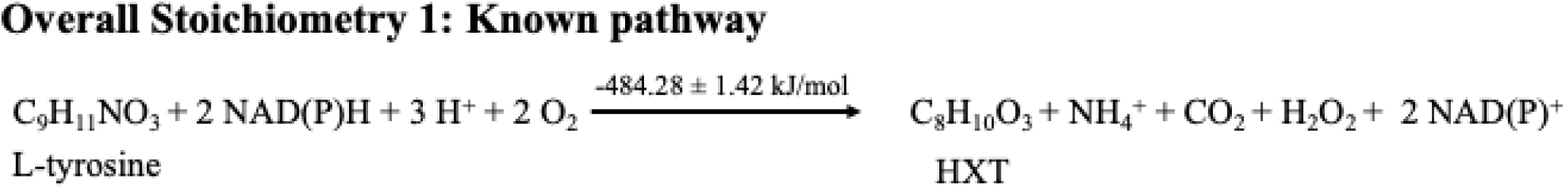

However, optStoic also identified the simpler 1 Tyr ➔ 1 HXT overall stoichiometry with the fewest co-products and co-substrates, and no cofactor usage:

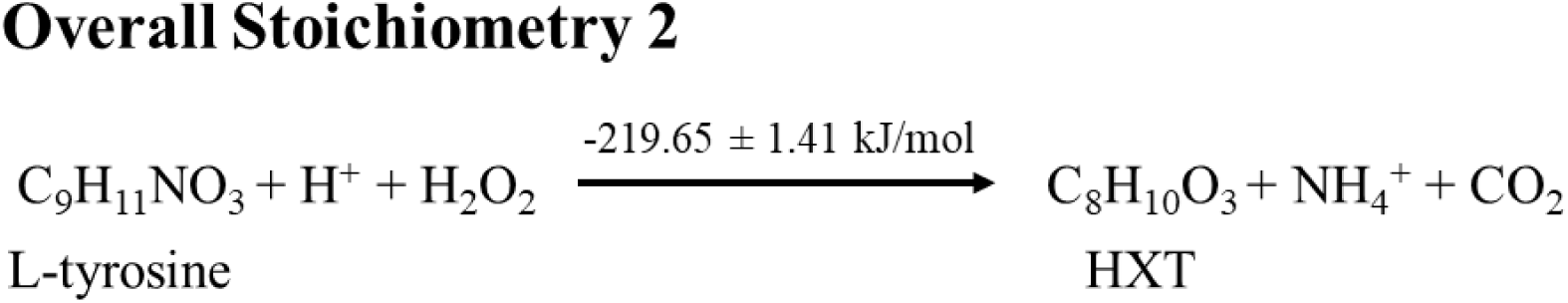

As shown above, both stoichiometries involve a high negative overall Gibbs energy change alluding to their thermodynamic feasibility.

The next step was to input these identified stoichiometries in novoStoic to construct complete pathways. Using stoichiometry 1, novoStoic found multiple pathways that have not been explored to our knowledge (see Supplementary Information File S1). Fig 7 illustrates these pathway alternatives, that encompass the existing pathway from the literature[12] alongside an alternate route.

**Fig 7:**
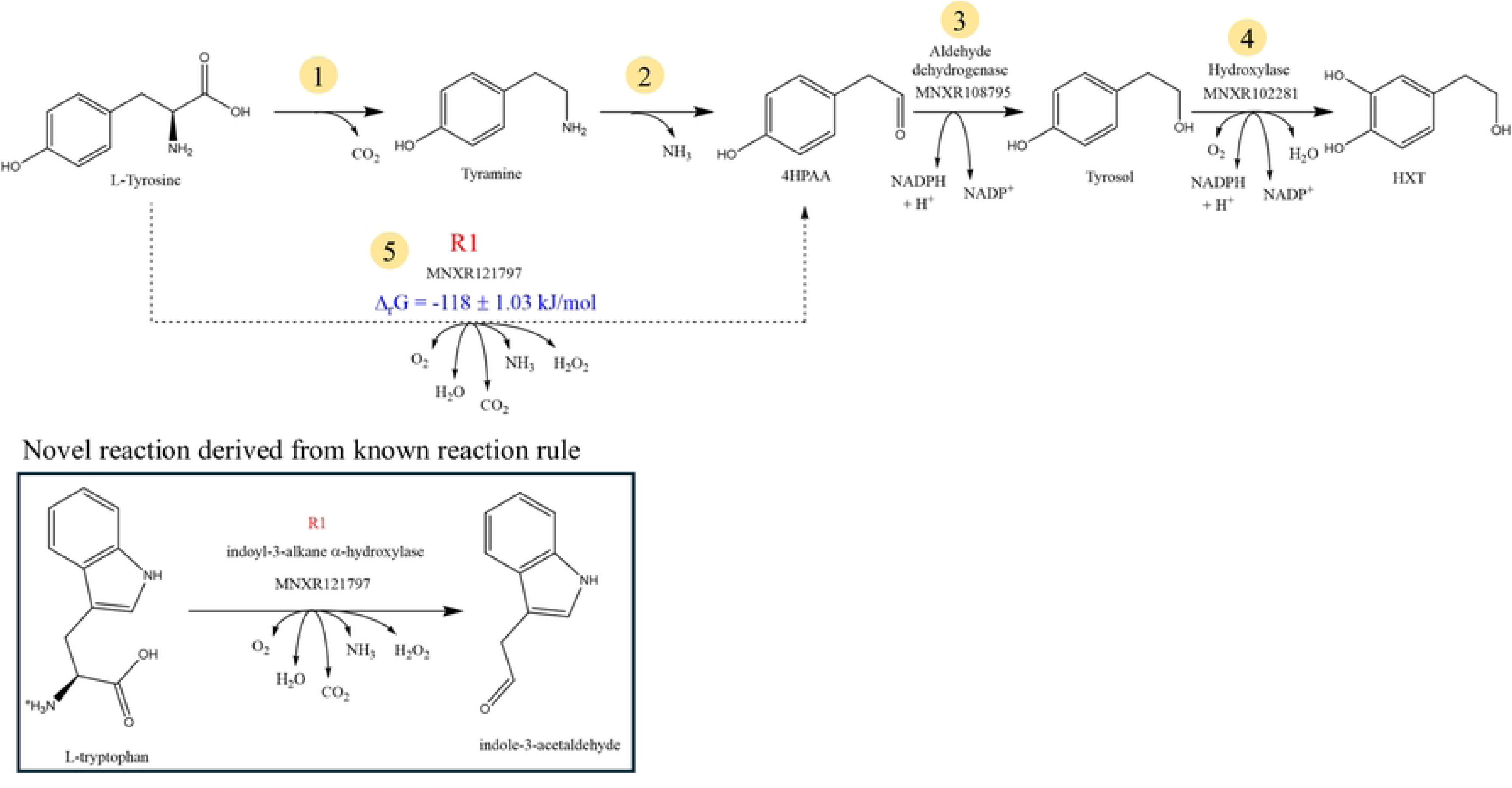
Pathway identified using novoStoic with overall stoichiometry 1. Here pathway with steps 1-4 shows the known pathway from the literature. Whereas the pathway with steps 5-3-4 shows a novel pathway that bypasses steps 1 and 2 in a single step. Here step 1 is a decarboxylation reaction, step 2 is tyramine: oxygen oxidoreductase, step 3 is aldehyde dehydrogenase, step 4 is a hydroxylase reaction and novel step 5 comes from a reaction MNXR121797 which is a hydroxylase enzyme.

The known pathway involves steps 1-4 (Fig 7) converts L-tyrosine to tyramine via tyrosine decarboxylase enzyme (EC 4.1.1.25) followed by the hydroxyl group addition at the meta-position to convert tyramine to 4HPAA using tyramine: oxygen oxidoreductase, followed by dehydrogenase and hydroxylase enzymes to convert 4HPAA to tyrosol and HXT, respectively. However, an alternative novel pathway was identified that bypasses the first two steps of converting tyrosine to 4HPAA by integrating them into a single step using the reaction rule R1 (i.e., EC 1.13.99.-) derived from reaction indoyl-3-alkane-α-hydroxylase. The pathway in Fig 7 shows the novel bypassing step 5 which follows the reaction rule of reaction MNXR121797. Using dGPredictor, we confirmed that the proposed novel step involves a negative standard Gibbs energy. Reaction MNXR121797 (EC 1.13.99.-) has a single enzyme sequence assigned to it which can serve as the starting enzyme that would need to be re-engineered for usher activity on the new substrate L-tyrosine. This makes the use of EnzRank redundant. Even though the reaction for step 4 is not cataloged in MetaNetX it follows the reaction rule of reaction MNXR102281. However, recent experimental evidence[12] points out that enzyme 4-hydroxyphenylacetate 3-monooxygenase has a secondary activity on tyrosol obviating the need for enzyme discovery/engineering.

Using optimal stoichiometry 2, we identified novel pathways using three steps that do not use any cofactors and only small molecules co-reactants and co-products (see Supplementary Information File S1). Fig 8 shows one of the pathways identified by novoStoic for the synthesis of HXT.

**Fig 8:**
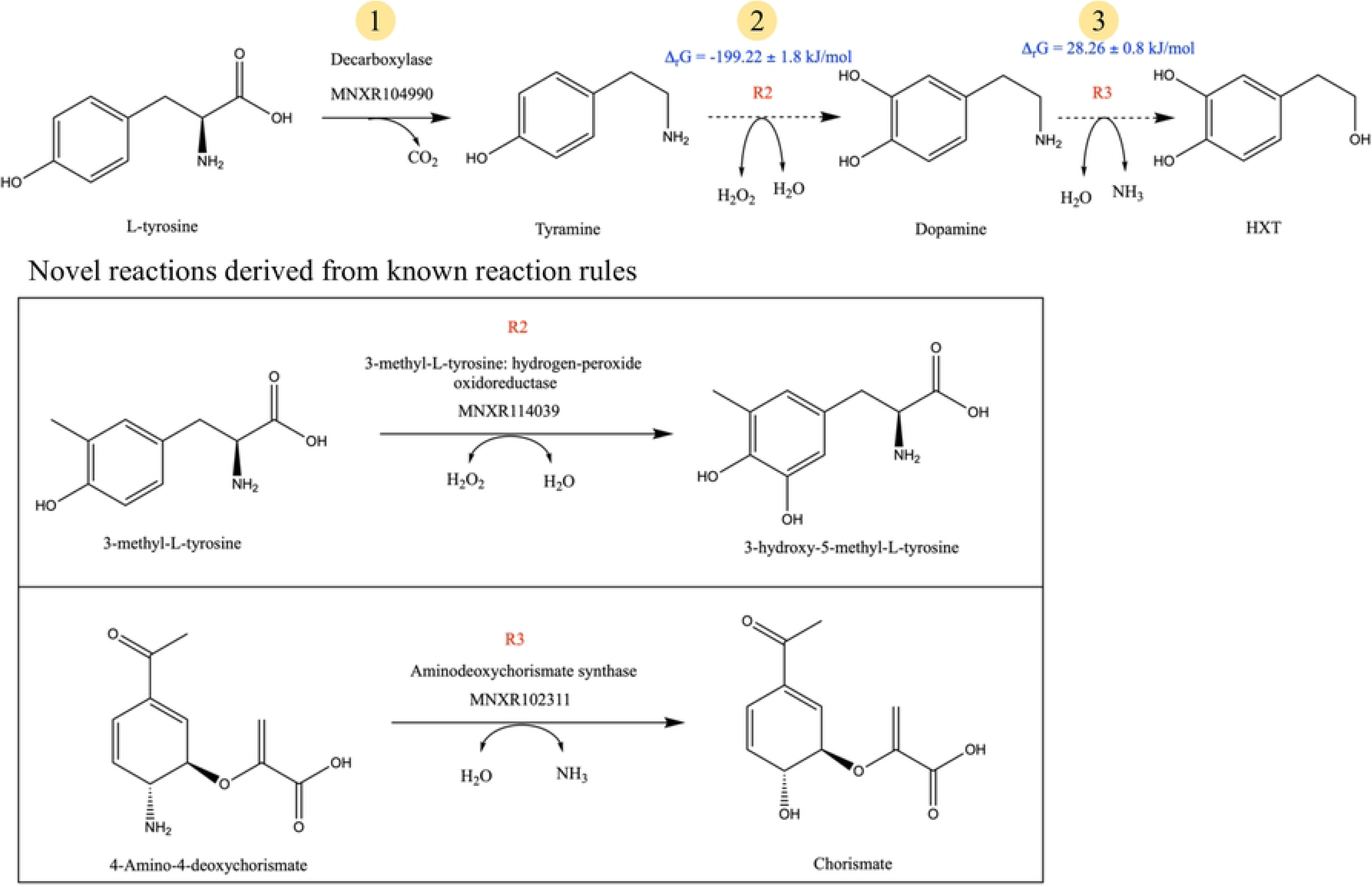
Pathway identified using novoStoic with overall stoichiometry 2. Here pathway with steps 1-3 shows the shorter novel pathway. Here, step 1 is a known decarboxylation reaction, step 2 is derived from the reaction rule R2, and step 3 is derived from reaction rule R3.

Fig 8 illustrates the three-step cofactor-free pathway. Step 1 is the same as in the known pathway shown in (Fig 7)[12]. The novel reaction associated with step 2 follows the reaction rule associated with reaction 3-methyl-L-tyrosine hydrogen peroxide oxidoreductase (EC 1.11.2.5) that performs a hydroxyl addition to convert tyramine to dopamine. Notably, there is a known reaction in the MetaNetX database that performs this conversion, but it requires NAD(P)H as a cofactor. Using dGPredictor, we assessed the thermodynamic feasibility of the desired direction and found a high negative standard Gibbs energy. Step 3 is a novel reaction that shares the same reaction rule as reaction MNXR102311 (aminodeoxychorismate synthase) which converts 4-amino-4-deoxychorismate to Chorismate. dGPredictor assesses the standard Gibbs energy change to be slightly positive, which can be made feasible by tilting the reactant concentration to move the reaction in the forward direction. Nevertheless, this example showcases that novoStoic can identify a shorter pathway with a lesser number of cofactors compared to the known pathway. The novel step R2 (Fig 8) was already suggested in the recent article by Chen et. al[12] that includes the engineering promiscuous enzyme 4-hydroxyphenylacetate 3-monooxygenase to show activity with tyramine. However, novoStoic qualified it as a novel step as the reaction was not present in the database yet. For the novel step R3 (in Fig 8), we found a total of 3,744 unique enzyme sequences from the reactions with the same rule as R3 (see Supplementary Data File S2 provides the list of all the enzymes for step R3). Next, EnzRank was used to rank-order known enzymes for the same reaction rule from which the novel step R3 is derived using the probability score to identify the potential of an enzyme to act on a substrate (dopamine). Here, Table 1 shows the top 3 candidates found using EnzRank for reaction step R3, and Supplementary Data File S3 contains the EnzRank scores for all the enzyme sequences for the same rule as R3.

**Table 1:**
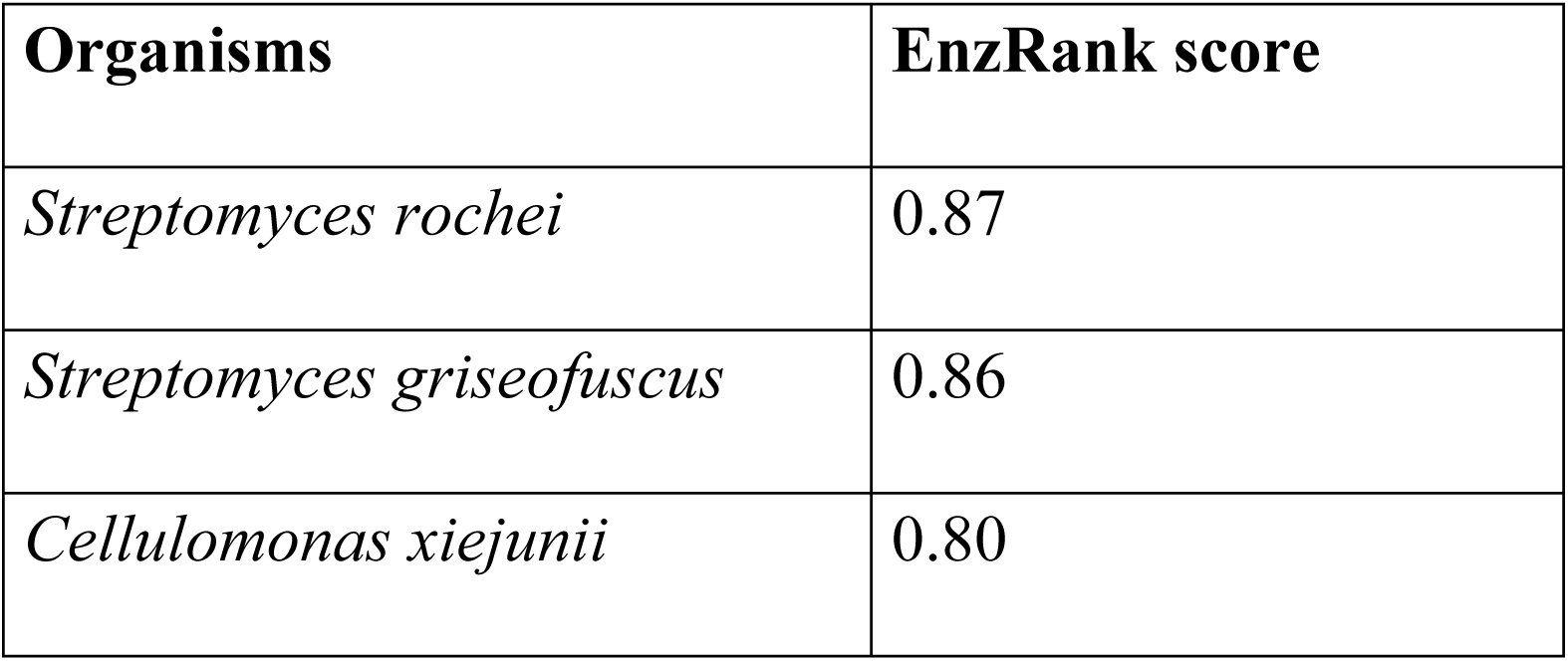
Top 3 genes from different organisms selected by EnzRank for novel reaction step R3 with reaction rule derived from aminodeoxychorismate synthase enzyme.

Using HXT as an example/case study, we highlighted how novoStoic2.0 can be used to find multiple pathway alternatives for HXT synthesis mapping to different overall stoichiometries. These alternatives mapping to different overall stoichiometries were subsequently subjected to thermodynamic feasibility analysis and enzymes were prioritized for all novel steps.

In collaboration with the Molecule Maker Lab Institute (MMLI) at UIUC, novoStoic2.0 is being developed as an interactive website for the synthesis planning tool will be a part of the AlphaSynthesis and will be made publicly available at http://novostoic.platform.moleculemaker.org/.

## Discussion

Pathway design entails multiple tasks ranging from decisions on starting points and possible co-reactants/products, sequence of metabolic reaction steps including novel reaction by-passes, selection of enzymes for uncharacterized reaction steps, checking on thermodynamic feasibility of chosen reaction direction, and many more. novoStoic2.0 integrates many of these steps within a single resource thus streamlining the task of pathway discovery and evaluation. By enabling the rapid computational exploration of design alternatives, it expands the space of alternatives explored and thus the chances of success. The development of an easy-to-use web-based interface consolidates many tasks into a single platform.

novoStoic2.0 is by no means inclusive of all tasks needed to instantiate a pathway design. The pathway will ultimately have to be ported in a production strain. The strain will have to be engineered so that no carbon flux leaks away from the desired pathway. The expression levels of genes and translation rates of proteins along the pathway will have to be finely tuned to limit metabolic burden and inhibitory controls will have to be ameliorated. There is already a rich literature of tools aimed at addressing these challenges (e.g., optKnock[32], optForce[33], RBSCalculator[34], etc.). Moving beyond, pathway carbon and energy efficiency additional design considerations are equally important. These may include predicting the toxicity of intermediates, safeguarding against protein misfolding/aggregation, presence of high-affinity product exporters, etc. For all these tasks several computational tools are available to provide estimates [35–38]. Finally, perhaps the most challenging step is the identification or redesign of enzymes with the desired substrate specificity and activity to carry out novel conversion steps. Whereas in some cases promiscuous enzymes can be found or adapted through directed evolution[12,39] rapid advances in enzyme design using ML tools promise to automatize this step[40–42]. We envision that many of these aforementioned tools will be integrated into future versions of novoStoic.

We anticipate that ML techniques will likely revolutionize pathway design in the same way that they have changed the landscape in protein folding and enzyme design. For instance, leveraging Large Language Models (LLMs) on up-to-date literature information could immediately inform pathway designs. For example, for the identified pathway shown in Fig 7, novoStoic classified step 3 converting tyrosol to HXT as a novel step but a recent article by Chen et. al[12] suggested that the desired novel activity has already been assigned an enzyme that was not present yet in the database. Therefore, LLMs offer the promise of automating data mining from the literature and moving beyond simple SMILES encodings for metabolites and reactions. novoStoic2.0 already leverages some of these ML developments and offers a versatile platform to integrate additional ones in the future.

## Methods

This section describes the design, implementation, and use of the novoStoic2.0 user interface. We used Streamlit-based Python framework to build the user interface for all the tools that are integrated within novoStoic2.0. We used the MetaNetX database to extract a total of 74,612 reactions and 1,292,153 molecules. Upon processing the database by removing unbalanced reactions, transport reactions, and reactions containing generic molecules, we have 23,585 reactions. Similarly, for molecules, we removed the generic molecules, molecules with multiple MetaNetX IDs, and used molecules that were present in 23,585 reactions, that ended up using 17,154 molecules from the database. Hence, in novoStoic2.0, we use 23,585 reactions and 17,154 molecules for pathway design. Using that, we generated the molecular signature for the molecules and reaction rules for the reactions that are used in novoStoic to design de novo pathways for biosynthesis. Finally, we generated 9,686 unique reaction rules from 23,585 reactions. However, both dGPredictor and EnzRank use the KEGG database for standard Gibbs energy estimations and enzyme selection, respectively. We created a mapping of MetaNetX IDs with KEGG IDs, in order to use the novoStoic outputs in both the tools. If a MetaNetX ID is not present in KEGG, we use InChI and SMILEs string and consider them as novel molecules in dGPredictor and EnzRank. Using dGPredictor to estimate the standard Gibbs energy change for novoStoic identified reactions, we generate thermodynamically feasible pathways. Allowing the integration with EnzRank, we probe all the enzyme sequences for a given reaction rule from the KEGG and Rhea database using the KEGG REST API[31] and Rhea API[43] along with the novel substrate SMILEs string to estimate the probability score for enzyme-substrate activity and rank-order the known enzymes for the novel reaction steps.

## Data and Code Availability

The data used in building the platform is available at ScholarSphere (https://doi.org/10.26207/fxd2-se27) and the source code for running the Streamlit version locally for the web interface is available on GitHub (https://github.com/maranasgroup/novoStoic2.0).

## Acknowledgements

This work is supported by the U.S. National Science Foundation funded Molecule Maker Lab Institute (MMLI), award number 2019897 supported by National AI Research Institutes Program of the Directorate for Computer and Information Science and Engineering (CISE), in collaboration with the Division of Chemistry (CHE) and the Division of Chemical, Bioengineering, and Environmental Transport Systems (CBET) awarded to CDM. The funders had no role in study design, data collection and analysis, decision to publish, or preparation of the manuscript.

I acknowledge the NCSA team at the University of Illinois Urbana-Champaign for their invaluable assistance in developing the novoStoic2.0 website, an integral component of the AlphaSynthesis platform for the Molecule Maker Lab Institute (MMLI). I extend special thanks to Matt Berry, Lijiang Fu, Kate Arneson, Bingji Guo, and Sara Lambert for their significant contributions.

## Contributions

V.U. and C.D.M. planned the work. Code for novoStoic2.0 was written by V.U., code for data extraction and optStoic was written by M.A., and tested pathway generated by V.U. Initial draft of the manuscript written by V.U., with subsequent contributions from M.A. and V.U. All authors have given approval to the final version of the manuscript.

## Supplementary Information

Supplementary Information, File S1

Supplementary Data, File S2

Supplementary Data, File S3

